# Variable Chlorhexidine Tolerance Across *Klebsiella* Species from a Single Facility

**DOI:** 10.1101/2025.09.22.677961

**Authors:** David Lehman, Aubrey E Hetzler, Tayloe Friedrich, Madeline M Strouse, Katie E Barry, Shireen M Kotay, Amy J Mathers

**Affiliations:** Division of Infectious Diseases & International Health. Department of Medicine, University of Virginia School of Medicine. Charlottesville, Virginia, USA; Clinical Microbiology, Department of Pathology, University of Virginia, School of Medicine, Charlottesville, Virginia

**Keywords:** Chlorhexidine tolerance, Hospital Environment, Carbapenemase Producing Enterobacterales, *Klebsiella quasipneumoniae*

## Abstract

We assessed in vitro chlorhexidine tolerance across *Klebsiella* species from a hospital with widespread chlorhexidine use. Isolates underwent MIC testing and whole genome sequencing. *Klebsiella* species showed varying tolerance, with *K. quasipneumoniae* having the highest MICs. No clear link to acquired resistance was found, though chromosomal efflux pumps appeared in some traditionally environmental species. Differences in chlorhexidine tolerance between species highlight the role that biocides could have in shaping microbial populations in the hospital environment.

---

The genus *Klebsiella* encompasses diverse species occupying both clinical and environmental niches. *K. pneumoniae sensu stricto* has emerged as one of the most consequential multidrug- resistant pathogens worldwide, with carbapenemase-producing *K. pneumoniae* representing an urgent public health threat and a major cause of hospital-associated infections in vulnerable patients(1). Hospital wastewater systems, including sink drains, P-traps, and toilets, are increasingly recognized as persistent reservoirs for carbapenemase producing Enterobacterales, and can allow a reservoir for carbapenemase gene exchange including among *Klebsiella* sp.(2, 3).

Species-level identification within the *K. pneumoniae* complex, including *K. pneumoniae sensu stricto, K. variicola*, and *K. quasipneumoniae*, is problematic using standard clinical microbiology methods and often requires whole-genome sequencing for accurate classification(4, 5). Although species across the *K. pneumoniae* complex have been documented to cause infections, their natural niches and virulence vary (6). Despite their prevalence in both patient and environmental reservoirs, the ecological and selective forces shaping species distribution within the hospital setting remain poorly defined, but carbapenemase gene exchange can occur between species(7, 8).

Chlorhexidine is widely used in healthcare as a patient skin antiseptic, hand hygiene agent, and equipment disinfectant(9, 10). While full resistance at clinically applied concentrations is unlikely for any Enterobacterales(11), interspecies variation in tolerance has been described, and selective pressure from antiseptic use is a longstanding concern in infection control(12, 13).

Outbreaks associated with *Serratia marcescens*, a species with intrinsic chlorhexidine tolerance, underscore the potential for disinfectants to influence species persistence and selection within hospital environments (11, 12). Whether such selective effects extend across the *Klebsiella* genus remains unclear, despite their importance as environmental and clinical reservoirs of antimicrobial resistance. This study leveraged whole-genome sequencing to accurately speciate *Klebsiella* isolates collected from environmental reservoirs and patients. Our goal was to evaluate whether chlorhexidine tolerance varies across the *Klebsiella* genus within a single hospital.

Isolates that were previously collected from the University of Virginia, where all intensive care unit hand hygiene soap is SCRUB-STAT (Ecolabs, St. Paul, MN) with 2% chlorhexidine gluconate. We took sequenced isolates across species and have had a historical focus on *bla*_KPC_- positive *Klebsiella* sp. isolates from patients and the environment between 2007 and 2024. We also used several known species from curated strains from the American Type Culture Collection (ATCC Manassas, VA) as species controls where available. Chlorhexidine was diluted in sterile water and then applied to Cation-Adjusted Müeller Hinton Broth (CAMHB), with final concentrations ranging from 0.125 μg/mL to 128 μg/mL in doubling dilutions using the methods described in Lutgring et al (14). To achieve the highest test concentration of 128 μg/mL without precipitation, an increase from 64 μg/mL previously described, both CAMHB and chlorhexidine solutions were pre-warmed to 37×C prior to combining. All isolates were tested in triplicate.

Initial speciation was done by MALDI-TOF (Biomerieux, Durham, NC) with further speciation and strain typing done via whole genome sequencing with paired end sequencing was done using a MiSeq v2 300 cycle Reagent Kit on the MiSeq (Illumina, San Diego, CA) platform. Species identification was performed by calculating the genomic distances of the read sets against a curated collection of complete bacterial chromosomal assemblies in National Center for Biotechnology Information (NCBI) RefSeq database using MASH(15)(v2.1). De Novo assembly was checked against the AMRFinder database in NCBI with >99% identity and coverage. Isolates were selected to capture a variety of species within the genus *Klebsiella* and to compare them with E. coli, which serves as a standard, and Serratia marcescens, known to exhibit higher MICs (19). To evaluate differences in log2-transformed MIC values among bacterial species, we first conducted a Kruskal-Wallis test to assess overall group differences (α=0.05). Post-hoc pairwise MICs between species with comparisons using Dunn’s test, implemented from the rstatix R package (v. 0.7.2), with a Bonferroni correction.

We tested 78 unique isolates; 57 clinical, 15 environmental and six ATCC against Chlorhexidine (Table 1). The technique had good reproducibility with all isolates being within one dilution thus 100% essential agreement across the triplicate testing with the median used as the MIC.

**Table 1.** Isolate Collection and Chlorhexidine MIC Testing Results.

|  | MIC50<br>µg/mL | MIC90<br>µg/mL | Range<br>µg/mL | Clinical | Standard | Environmental |
| --- | --- | --- | --- | --- | --- | --- |
| *Klebsiella pneumoniae*<br>sensu stricto<br>n=13 | 16 | 32 | 8-32 | 10 | 3 | 0 |
| *Klebsiella quasipneumoniae*<br>n=22 | 32 | 64 | 16-64 | 10 | 1 | 11 |
| *Klebsiella oxytoca*<br>n=3 | 8 | 16 | 4-16 | 2 | 0 | 1 |
| *Klebsiella aerogenes*<br>n=6 | 32 | 32 | 16-32 | 6 | 0 | 0 |
| *Klebsiella grimondii*<br>n=1 | - | - | 32-32 | 0 | 0 | 1 |
| *Klebsiella variicola*<br>n=5 | 32 | 32 | 16-32 | 5 | 0 | 0 |
| *Klebsiella michiganensis*<br>n=4 | 16 | 32 | 4-32 | 3 | 0 | 1 |
| *Escherichia coli*<br>n=9 | 2 | 8 | 1-32 | 7 | 1 | 1 |
| *Serratia marcescens*<br>n=15 | 128 | 128 | 32-128 | 14 | 1 | 0 |

In our study like others, *E. coli* generally exhibited lower MICs compared to all *Klebsiella* species including *K. pneumoniae* (p=0.02), *K. aerogenes* (p=0.001), *K. quasipneumoniae, K. variicola, S. marcescens* (all p<0.0001) but not *K. grimondii, K. oxytoca* and *K. michiganensis* although numbers are small (Fig 1). A larger prior evaluation showed *E. coli* had an MIC_90_ of 16 µg/mL whereas, *K. pneumoniae* demonstrated an MIC_90_ of 32 µg/mL(20). Interestingly, in this collection, several *K. quasipneumoniae* isolates, many which were derived from environmental sources, displayed higher MICs, with an MIC_50_ of 32 and MIC_90_ of 64 µg/mL. Using the Dunn statistical test MICs of *K. quasipneumoniae* were significantly higher than *K. oxytoca* (p=0.002), *K. pneumoniae sensu stricto* (p=0.004).

**Figure 1.**
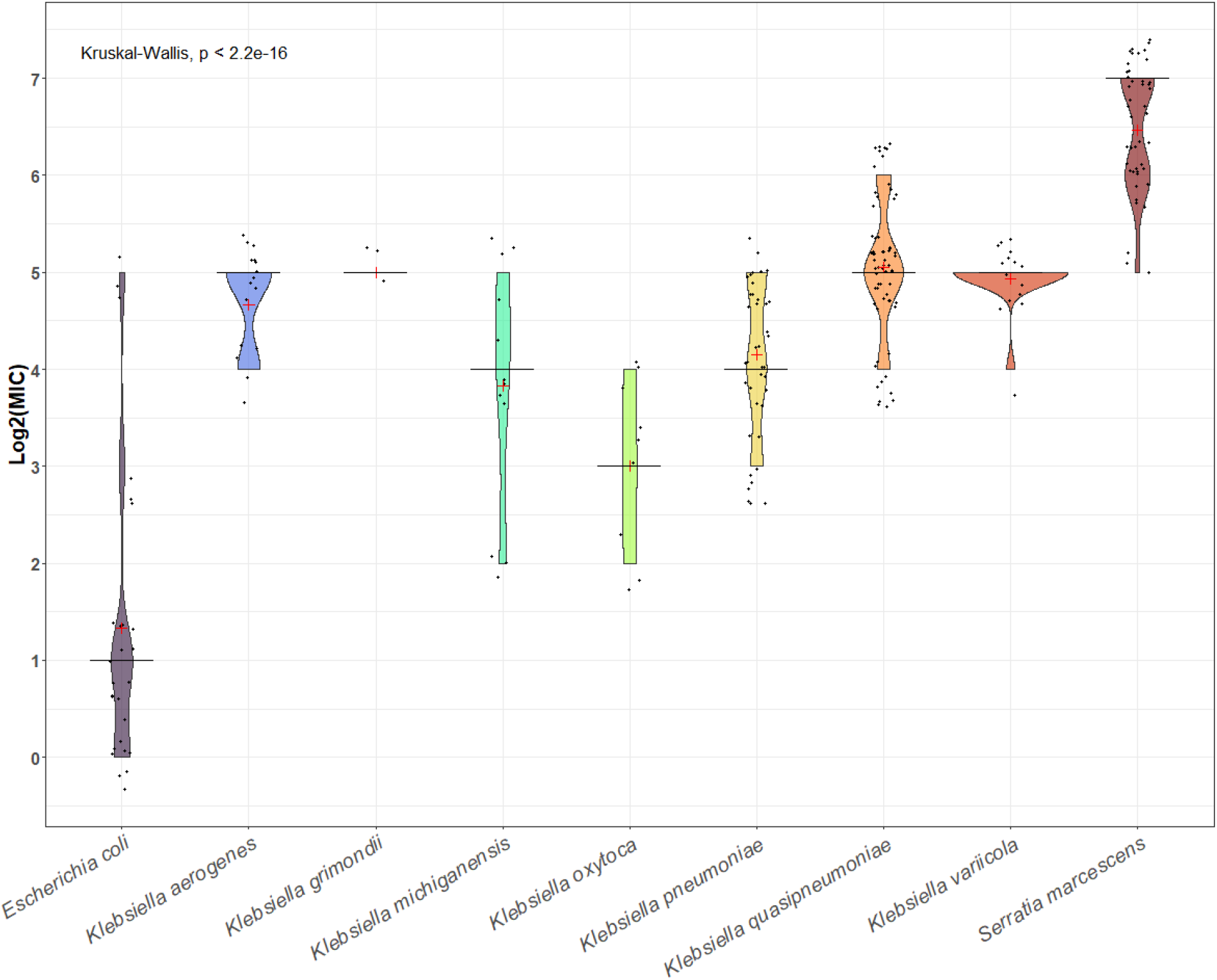
Violin Plots depicting MIC distribution for chlorhexidine by species. Red plus signs indicate the mean log_2_(MIC) concentration for each species, while black horizontal bars represent the median values. A Kruskal-Wallis test indicates an overall statistically significant difference in MIC distributions across species.

We further evaluated resistance gene content across this multidrug-resistant collection using AMRFinder (NCBI). Predicted genes included quaternary ammonium compounds (qac) determinants that may be plasmid-borne and drive transferrable efflux in gram positives but these were not consistently found nor was there a clear correlation with chlorhexidine MICs (Supplemental Fig 1 and Fig 2.). Previous studies in gram negative bacilli have failed to demonstrate that mobile AMR genes promote chlorhexidine tolerance conclusively but rather the resistance is more like selected by chromosomal genes and mechanisms allowing tolerance such as outer membrane differences and effective efflux pumps (23, 24). Interestingly both *the K. variicola, K. quasipneumoniae* consistently had resistance nodulation cell division (RND) family oqxB efflux pumps as did several of the *K. pneumoniae* (25).

At our institution, where chlorhexidine-impregnated soap is routinely used, we have found ongoing recovery of *bla*_KPC_-positive isolates from hospital wastewater plumbing, particularly sink drains(21). Sequencing has revealed that several isolates initially identified as *K. pneumoniae* were further speciated via sequencing as *K. quasipneumoniae* (8). We have also frequently detected *bla*_KPC_-positive *Serratia marcescens* in this plumbing reservoir and a strain that has been seen in patients and the environment within our hospital was included (CAV1492)(22).

These preliminary findings suggest a potential link between environmental persistence, biocide tolerance, and antimicrobial resistance. However, the small sample size and the selection from a single institution means the data should be viewed as very preliminary. Species such as *K*.

*variicola* and *K. quasipneumoniae*, which are more commonly associated with environmental reservoirs, showed generally higher minimum inhibitory concentrations (MICs) for chlorhexidine. This raises the possibility that biocide exposure in healthcare or environmental settings selecting for strains with both increased tolerance to disinfectants and the ability to carry clinically relevant resistance determinants such as *bla*_KPC_ driving outbreaks related to premises plumbing (26, 27).

Although the dataset is limited in size, it highlights the importance of species-level identification and emphasizes the need to consider both clinical and environmental isolates in infection control strategies. The observed differences in chlorhexidine tolerance between species underscore the role of ecological pressures in shaping microbial populations. Co-selection of antimicrobial resistance through biocide exposure may be an underappreciated factor in the emergence and spread of multidrug-resistant organisms in healthcare settings, warranting further investigation into the mechanisms of tolerance and their epidemiological consequences.

## Acknowledgments

Support for the work and DL effort on the project was funded by the National Institute of Health (Bethesda, MD, USA) training grant 5T32AI007046.

## Author Contributions

David Lehman did much of the experimental work for susceptibility testing, analysis and writing of the original manuscript draft, Aubrey Hetzler oversaw with laboratory experiments and visualization of the data as well as analysis and editing, Tayloe Friedrich performed laboratory experiments, Madeline M Strouse performed laboratory experiments, Katie Barry performed whole genome sequencing and oversaw laboratory work, Shireen Kotay assisted with writing, visualization and genomic data analysis, Amy J Mathers conceptualization, investigation, project administration, supervision of writing of original draft, review and editing

## Data Availability

All sequenced data generated and analyzed in this study were deposited at NCBI under BioProject ID PRJNA948355.

## Ethics Approval

All clinical isolates were collected as discarded isolates with the review and approval of the University of Virginia IRB#13558.

## Additional Files

The following material is available online.

Figure S1. AMR Gene profiles for each isolate of *Klebsiella* sp.

Figure S2. AMR Gene profiles for each isolate of *E. coli and S. marcescens*

